# A pilot study for whole proteome tagging in *C. elegans*

**DOI:** 10.64898/2026.02.09.704846

**Authors:** Matthew Eroglu, Oliver Hobert

## Abstract

Tagging all proteins encoded by an animal genome with a fluorescent tag would open many windows to the discovery of unexpected patterns of protein expression and localization. To scale such an approach, it would be beneficial to introduce multiple, spectrally distinct fluorophore tags in parallel. As proof of concept for scalable pooled tagging, we undertook a pilot study in the nematode *C. elegans,* in which we set out to tag 30 different genetic loci with three different fluorophores, with 3 tags being introduced at a time. By choosing essential genes, predicted based on transcriptomics to cover a range of expression levels, we explore issues relating to disrupting gene function and visibility of tagged proteins. We demonstrate that such a tagging approach is highly efficient and indeed reveals unanticipated patterns of cellular sites of expression, as well as subcellular protein localization. We hope that this pilot study will motivate attempts to scale this tagging approach to more loci and, ultimately, the whole genome.

## Introduction

Gene transcript-based expression atlases have provided invaluable insights into animal biology. The next frontier lies in generating whole animal protein expression atlases with single cell resolution that capture the cellular and subcellular specificity of protein expression and localization and visualize the dynamics of these parameters during development and in response to changes in internal and external states. The establishment of protein expression atlases is particularly urgent in light of the generally appreciated discordance between transcript detection – often used to infer cellular functions – and protein expression, a reflection of multifarious layers of posttranscriptional gene regulation (Liu et al., 2016; Schwanhäusser et al., 2011; Vogel and Marcotte, 2012).

One means to acquire protein expression and localization on a single cell level with whole proteome coverage lies in tagging all genes in the genome with a fluorescent reporter. This approach has been pioneered in yeast (Huh et al., 2003), where chromosomally GFP-tagged strains provided the first cellular map covering ∼75% of the yeast proteome, assigning localization for 70% of previously unlocalized proteins while uncovering the compartmental logic underlying protein function and interactions. Tagging proteins in multicellular organisms provides a host of new insights since the differentiation of cells into distinct types and their assembly into tissues deploys many more proteins and encompasses the establishment and dynamic control of more complex protein expression and localization patterns.

We envision the tagging of the entire proteome of a multicellular organism to be most feasible in the nematode *C. elegans*. CRISPR/Cas9-mediated insertion of fluorophore tags (Dickinson et al., 2013) is now routinely used to visualize protein expression and localization in *C. elegans*. With procedural advances including direct injection of Cas9 ribonucleoprotein (RNP) complexes (Paix et al., 2015) and repair template preparation (Eroglu et al., 2023; Ghanta and Mello, 2020), such engineering is highly effective and rapid, given the short generation time of *C. elegans*. The transparency of the organism facilitates imaging of fluorophores in live animals. Advances in light-sheet fluorescence microscopy, such as dual-view inverted selective plane illumination microscopy (diSPIM), have enabled high-resolution, 3-dimensional live imaging of dynamic fluorophore localization patterns throughout all stages of development (Kumar et al., 2014; Wu et al., 2017). Super-resolution imaging approaches such as image scanning microscopy (ISM) (Huff, 2015) and stimulated emission depletion (STED) microscopy (Rankin et al., 2011) enable increasingly finer localization of proteins in live worms. Further resolution can also be achieved in fixed worms with expansion microscopy, without the requirement for specialized microscopy equipment (Yu et al., 2022). Extensive anatomical atlases are either already available (Altun et al., 2002) or in the process of being generated (Cook et al., 2019; Santella et al., 2022; Witvliet et al., 2021), allowing the mapping of fluorophore localization patterns onto anatomical features at electron microscopical scales (Vergara et al., 2021).

There is also ample precedent in the *C. elegans* literature that reporter tagging results in unanticipated insights. To cite two recent examples from our own lab, we found that Golgi-resident proteins displayed cell type-specific expression patterns not anticipated by transcriptomic analysis (Leyva-Díaz et al., 2025). As another example, the subcellular localization pattern of a DEG/ENaC channel in enteric neurons indicated unexpected functional insights into the protein (Bayer et al., 2025). It is also notable that the widely appreciated discordance of mRNA and protein expression levels is evident in *C. elegans* (Grün et al., 2014; Reilly et al., 2022, 2020), attesting to the need to extend currently existing transcriptome atlases (Gao et al., 2024; Ghaddar et al., 2023; Packer et al., 2019; Roux et al., 2023; Taylor et al., 2025, 2021) with protein expression atlases. Individual labs have also begun to tag entire protein families, such as extracellular matrix proteins, yielding novel insights into tissue organization and differential localization patterns of proteins within the same class (Ragle et al., 2025).

Currently, around 1554 proteins representing 8% of the proteome are estimated to have been endogenously tagged (Leyhr et al., 2025). However, at current rates with individual labs tagging specific proteins of interest without coordination, covering the proteome is projected to take around 100 years and, as past experience has shown, likely involve numerous duplicate attempts on a small number of commonly studied proteins (Leyhr et al., 2025). It would thus be beneficial for the field (a) to coordinate tagging efforts and (b) to scale up tagging protocols to enable coverage of the entire genome at a reasonable timescale and cost. Since the number of injections is a major time-limiting factor, pooling multiple injections into one would at minimum cut tagging time by a factor of 3. In *C. elegans*, screening for novel CRISPR/Cas9-induced genomic edits is already facilitated either by use of co-injection markers (i.e., plasmids that form extrachromosomal arrays) that yield phenotypes or fluorescence in progeny of successfully injected worms, or co-editing well characterized loci using established and highly efficient reagents which likewise yield visible phenotypes. In the latter approach, termed “co-CRISPR”, worms edited at the marker locus are most likely to also carry the intended edit (Arribere et al., 2014); edits at F1, often being heterozygous, enable independent segregation of coCRISPR targets. The current efficiency of CRISPR/Cas9 mediated genomic editing enables simultaneously insertion of multiple fluorophores (e.g., mNeonGreen and mScarlet) as well as a co-CRISPR marker (*dpy-10*) at three independent loci in a single injection (Eroglu et al., 2023; Paix et al., 2015). These attempts pooled reagents previously established to work efficiently and targeted genes that were known to yield functional fusion proteins when tagged. Thus, while in principle current methods could allow tagging of at least 3 independent loci in one injection if a co-CRISPR marker is omitted, it is not known to what extent such an approach could be generalized across the genome with previously unvalidated reagents (i.e., guides and repair template homology arms) at novel loci to yield functional tags.

As proof of concept for scaling such a CRISPR/Cas9-engineered project to the whole genome, we report here the results of a pilot study in which we aimed to CRISPR/Cas9-engineer reporter tags into 30 genomic loci, generated through simultaneous tagging of three genes at a time. A subset of these 30 genes reveals unexpected patterns of expression and localization, illustrating how protein tagging approaches provide a route for new discoveries. We discuss here several aspects of such tagging approaches, highlighting promises, as well as limitations.

## Results

### Pipeline for pooled high throughput tagging by CRISPR/Cas9

Endogenously tagged genes can be detected by fluorescence stereomicroscopy if the fluorophores are expressed at sufficiently high levels and hence are readily visible as heterozygotes in F1 by visual screening without co-CRISPR markers. We reasoned that ubiquitously and highly expressed genes could be used as insertion markers and pooled with low or modestly expressed genes, with a first round of visual screening of the highly expressed genes followed by targeted PCR or higher power imaging of the lower expressed genes (**Fig. 1A**), thus eliminating the need for additional co-CRISPR markers.

**Figure 1:**
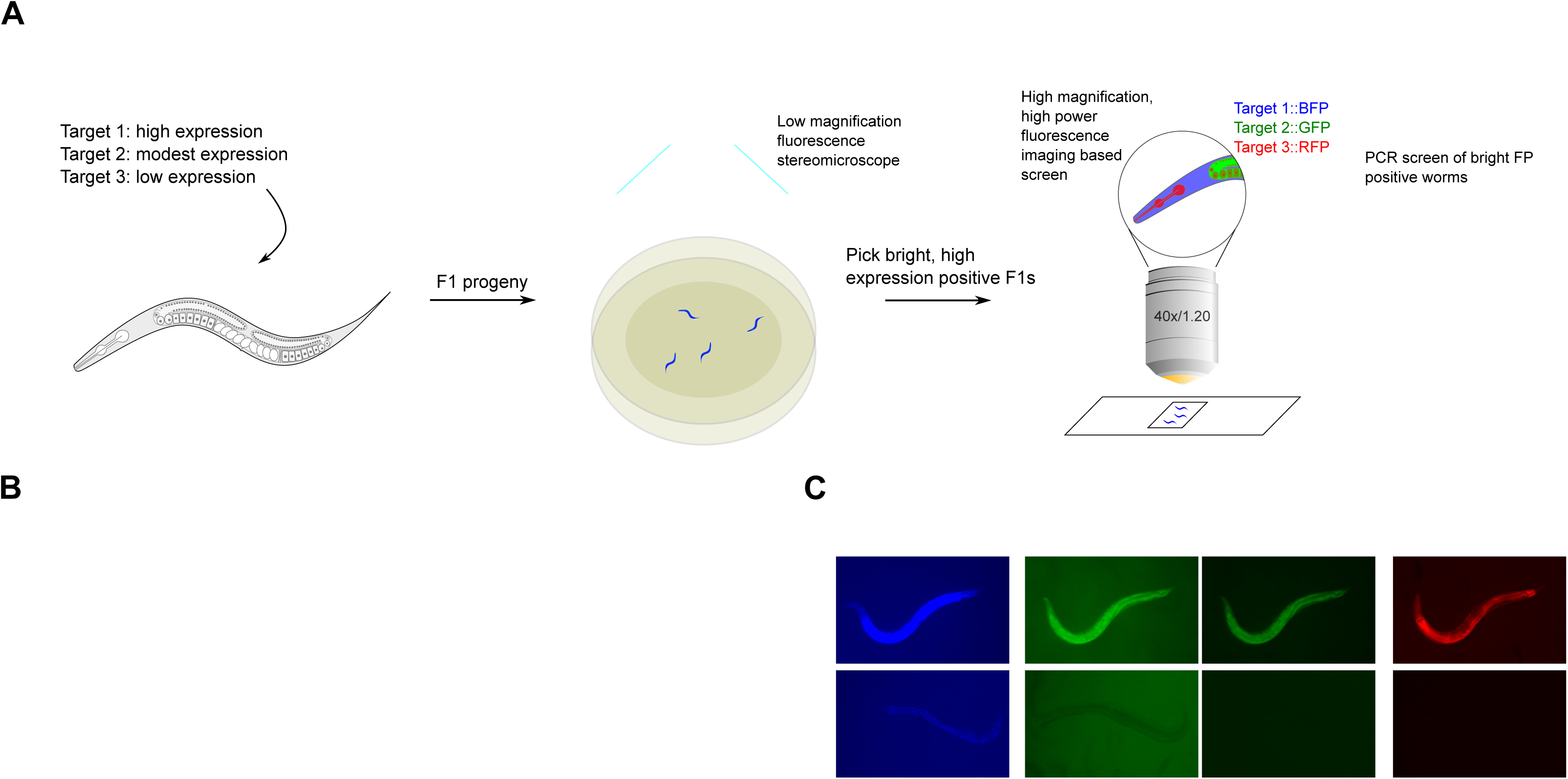
Pipeline for triple pooled CRISPR-Cas9 mediated gene tagging. **(A)** Schematic of injection and screening strategies. FP, fluorescent protein. **(B)** Excitation and emission spectra of selected fluorescent proteins used in the pilot study. Theoretical brightness of the fluorophores, calculated as a product of the extinction coefficient and quantum yield, is annotated as numbers adjacent to the respective emission spectra. The spectra and brightness values were adapted from FPbase (Lambert, 2019). **(C)** Practical visibility of fluorophores observed under a fluorescent stereomicroscope with different filter sets in live worms on NGM plates. Top, widefield images of worms tagged with the indicated fluorophore (FP) imaged through the indicated filter contrasted to non-tagged worms and background. Bottom, schematic illustration of observed view by eye under fluorescent stereomicroscope during screening. mRNA expression of the tagged protein in young adult worms is indicated in FPKM.

To facilitate screening by fluorescence stereomicroscopy and co-imaging, we chose the three currently brightest fluorescent proteins in the blue, green and red emission spectra: mTagBFP2 (Subach et al., 2011), mStayGold(J) (Ando et al., 2024), and mScarlet3 (Gadella et al., 2023), respectively (**Fig. 1B**). These tags are significantly brighter than first generation fluorophores, thereby allowing the detection of even very lowly expressed proteins. Autofluorescence in *C. elegans* is highest in blue and green wavelengths, and lowest in red. Furthermore, we noted that nematode growth medium (NGM) plates seeded with OP50 also contribute to background fluorescence with conventional blue and green filters on a fluorescent stereomicroscope (**Fig. 1C**). In contrast, we observed minimal background with yellow and red emission filters from worms as well as bacteria. We also noted from existing tagged strains that mStayGold could be observed brightly with YFP filters, which, combined with the minimal background, yielded a greater contrast with untagged worms enabling easier screening of mStayGold positive worms compared to conventional GFP filters (**Fig. 1C**). However, given the suboptimal excitation and emission of visualizing mStayGold with a YFP filter, mScarlet3 in practice yielded the greatest signal-to-noise ratio for visual screening. Thus, to balance fluorescence signal-to-noise ratios with predicted gene expression, we assigned the highest expressed genes to be tagged with mTagBFP2, modestly expressed genes with mStayGold, and lowest expressed genes with mScarlet3.

In our pilot experiments, we sought to establish three major principles: (1) the endogenous expression ranges at which fluorophores could be utilized for visual screening; (2) success rate of triple pooled tagging using novel guides and targets; (3) functionality of tagged proteins. We thus selected 30 genes across a variety of bulk transcript expression ranges which are generally predicted to be broadly expressed based on molecular function or, where molecular function is unknown (e.g., *ZK632.9*), single cell RNA sequencing (scRNA-seq) data (**Table 1**, **Fig. 2A, B**) (Gao et al., 2024; Ghaddar et al., 2023; Taylor et al., 2021). These proteins cover a wide range of predicted cellular functions and subcellular localization patterns (nuclear, cytosolic, transmembrane). 26 of 30 genes are annotated to display lethal, sterile, or other phenotypes by RNAi, enabling the assessment of tagged protein functionality.

**Figure 2:**
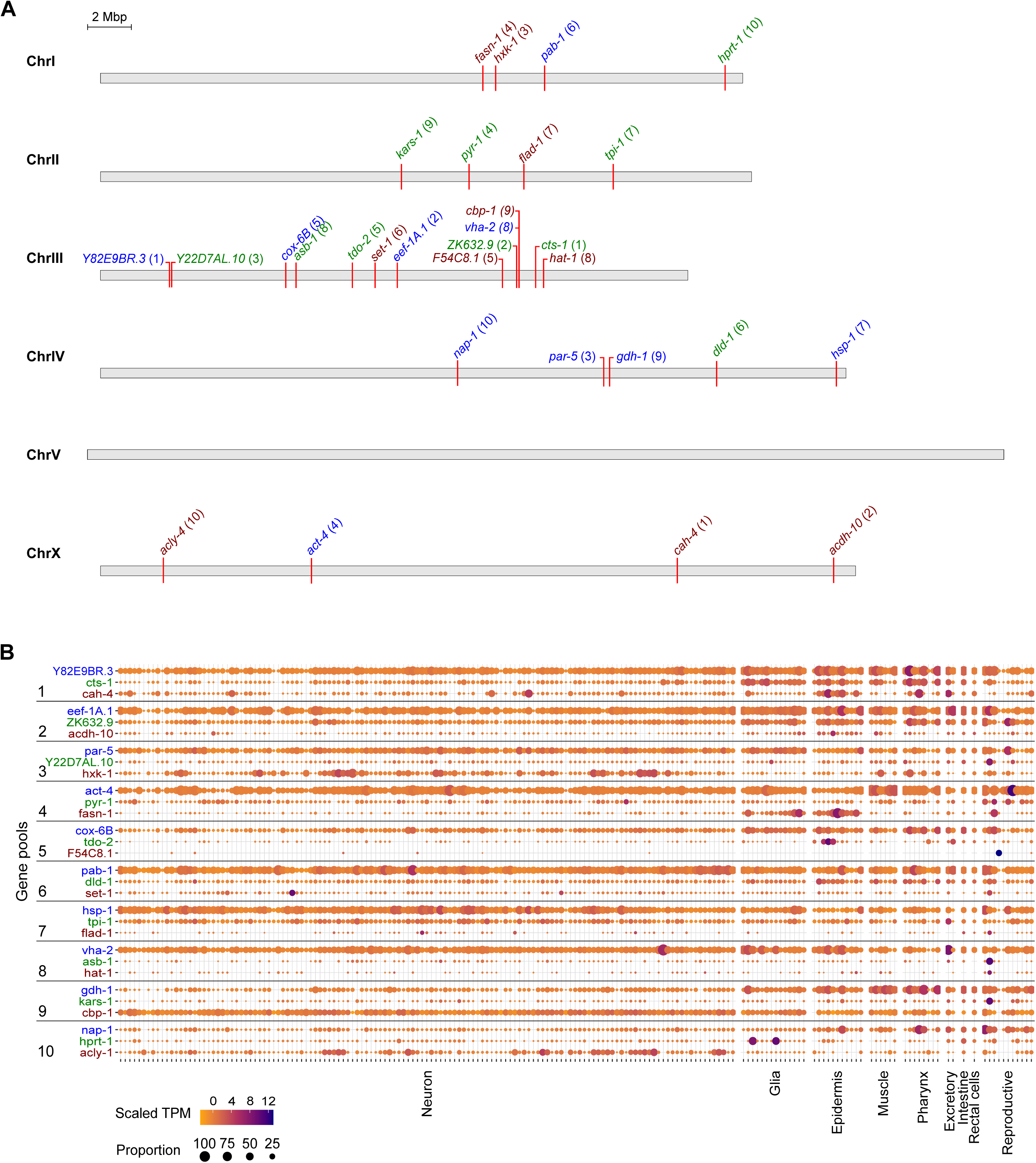
Selection of genes and predicted transcript expressions across tissues. **(A)** Location of selected genes on *C. elegans* chromosomes. Numbers in parentheses indicate the injection pool the gene was assigned to. Color of text indicates the fluorophore assigned to the gene: blue, mTagBFP2; green, mStayGold; red, mScarlet3. **(B)** Transcript expression levels of the selected genes across major tissues of *C. elegans* as measured by single cell RNA sequencing of adult hermaphrodites. Data was adapted from the CeNGEN adult hermaphrodite set (Taylor et al., 2021).

The *C. elegans* genome encodes around 20,000 protein-coding genes (Spieth et al., 2014). Approximately 6,700 marker genes would need to be sufficiently highly expressed to be used as visible markers to group with the entire genome in sets of three. In bulk RNA-seq of N2 worms at L4 stage, 6,804 genes are expressed above 20 fragments per kilobase of transcript per million mapped reads (FPKM) (Eroglu et al., 2024; Gerstein et al., 2010; Harris et al., 2020). Based on our prior experience comparing bulk transcript FPKM with visibility, we divided these genes into three subsets to assess the detection limit of the three fluorophores: ≥500 FPKM for mTagBFP2; 100-500 FPKM for mStayGold; and 18-100 FPKM for mScarlet3.

We tagged genes either at their N- or C-terminus, primarily aiming to capture as many predicted isoforms as possible while retaining a potential guide RNA target sequence within 10 nucleotides of the start or stop codon. Where both termini satisfied these conditions, we selected the C-terminus over the N-terminus, as previous studies have suggested that C-terminal tags are less likely to disrupt protein localization and expression (Davidi et al., 2019; Palmer and Freeman, 2004) and that genome wide tagging attempts in yeast have successfully used C-termini (Huh et al., 2003) with high concordance to prior annotated data. However, we acknowledge that subclasses of proteins such as peroxisomal proteins, C-tail- or glycophosphatidylinositol-anchored proteins, lipidated proteins and proteins with retrieval motifs, may only show correct localization when tagged at the N-terminus (Yofe et al., 2016).

### Efficient tagging across loci, expression levels and target pools

We co-injected the 30 selected targets as 10 pools (**Table 1**), with each pool being injected into 5 worms. Overall, we successfully isolated 8/10 mTagBFP2 tags, 9/10 mStayGold tags, and 7/10 mScarlet3 tags (**Fig. 3A, B**). All but one tag (*cox-6B*::mTagBFP2) was visible in the F1 generation of injected P0 animals, and these were subsequently isolated among F2 worms positive for the other tags in the pool. Fluorescent signals of HAT-1::mScarlet3 and CBP-1::mScarlet3 in the F1 progeny were dim but still sufficiently visible for quantification of knock-in efficiency, indicating that they are at the lower end of detectability for mScarlet3. In 4 pools, we isolated all 3 tags within the same worm, and in an additional experiment all 3 tags but in pairs. The remaining tags were isolated as pairs or singles, and all tags could also be isolated individually or segregated separately from heterozygous animals. Even when one or two tags in the pool failed, the remaining tags in the same pool could be independently isolated. Overall, 24 of 30 tags were successfully isolated, all of which were visible with fluorescence stereomicroscopy.

**Figure 3:**
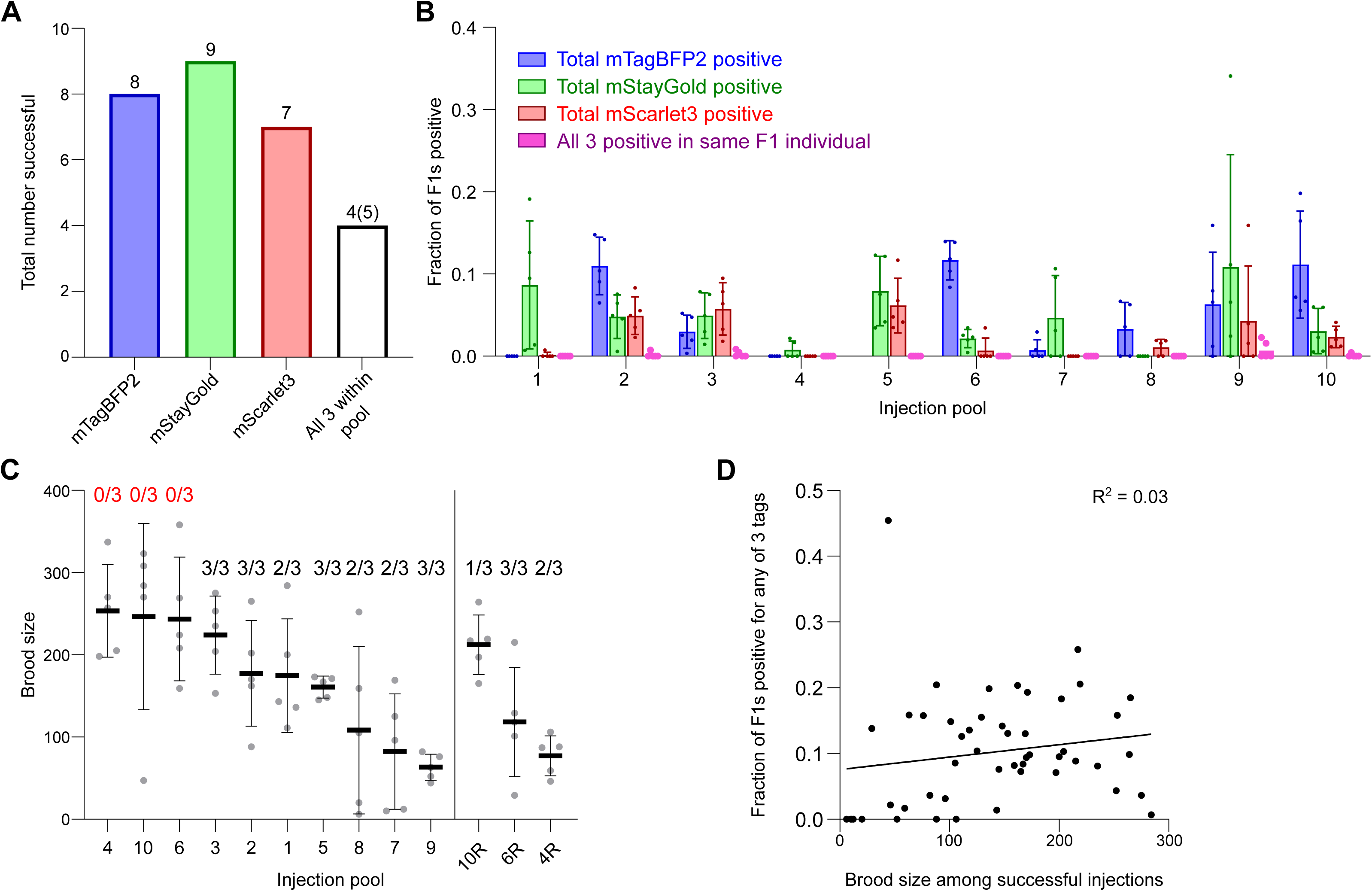
Successful tagging across loci, fluorophores and gene pools. **(A)** Total number of targets successfully tagged. White bar denotes all 3 genes within the pool were successfully tagged in any combination. In four pools, all 3 targets were isolated in a single, triple-tagged individual worm. In one pool, we obtained all 3 targets but in doubles. **(B)** Total insertion efficiencies per locus for the indicated targets across injection pools. Dots, fraction of F1s positive for indicated fluorophore per injected P0. Five worms were injected per pool except for pools 4, 10 and 6 which were repeated after initial failures as shown in panel C. Data shown in panel B only represents the repeat runs and omits the original failed injections. For group 5 including *cox-6B::mTagBFP2*, worms with detectable levels of mTagBFP2 fluorescence were not recovered in the F1 generation but were isolated among progeny of F1s positive for mStayGold and mScarlet3; we were thus unable to quantify efficiency for this locus at F1. Bars, mean ± SD. **(C)** Viable brood size of injected P0 worms represented as individual points. Lines, mean ± SD. Numbers above plots, number of successfully isolated tags out of 3 within the pool. 4R, 6R and 10R are re-injections of groups 4, 6 and 10. **(D)** Correlation of viable brood of P0s with fraction of F1 worms positive for any fluorophore tags. Goodness of fit is represented as an R^2^ value from a simple linear regression. Slope was not significantly greater than zero (P = 0.2497, F-test on Prism).

During our screening, we made several observations which enabled us to assess whether failures were due to technical or biological reasons. Our first attempt with pools 4, 6 and 10 yielded no positive progeny for any of the 3 targets in the respective pools despite the injected worms showing a healthy number of viable offspring (**Fig. 3C**). We reasoned that, given overall efficiency for individual tags, this was due to technical failures rather than the impossibility of functionally tagging all three genes within the pool. Indeed, upon reconstitution of the injection mixes with additional quality control steps (i.e., ensuring that repair templates were of sufficient purity as assessed by proper A260/230 and A260/280 ratios; see Methods), we successfully obtained 6/9 of the originally failed tags (**Fig. 3C**). In contrast, failures in obtaining one or two of the tags in the pool were generally interpreted as loss of function tags, and in these cases, we frequently observed fluorescent but dead embryos on the plates. While the combination of high brood size with 0/3 successful tags was a good indicator to reattempt individual pools, overall brood size across successful injections did not correlate with efficiency of insertion (**Fig. 3D**).

### Unanticipated cell type specific enrichment of targets

We assessed the expression patterns of the protein fusions by confocal microscopy in adult worms and compared them to expected tissue type specific abundances predicted by stage-matched scRNA-seq datasets (Gao et al., 2024; Ghaddar et al., 2023; Taylor et al., 2021). This revealed notable examples of mismatch between expected and observed expression patterns. For instance, the translation elongation factor EEF-1A.1 showed strong germline enrichment whereas it was predicted to be enriched in various cell types including germ cells, epidermis, intestinal and excretory cells by scRNA seq (**Fig. 4**). The acyl-CoA dehydrogenase ACDH-10 was somatically broadly and evenly expressed but strongly depleted in the germline, while we expected enrichment in intestinal, muscle and epidermal tissues given its role in fatty acid metabolism and transcript expression in scRNA-seq datasets (**Fig. 4**). Strikingly, the glucokinase HXK-1 showed strong enrichment in the gonadal sheath, whereas we expected ubiquitous expression with potential enrichment in glycolysis upregulated tissues such as muscle and neurons (**Fig. 5**). The lysyl-tRNA synthetase KARS-1 showed germline enrichment, while it was expected to be evenly distributed across tissues given its role in translation, but was correctly predicted by scRNA-seq to be germline enriched (**Fig. 6A-C**). The nucleosome assembly protein NAP-1 was most highly expressed in the germline, whereas it was predicted to be highest in the distal tip cell and pharynx (**Fig. 7A**). In contrast, the phosphoribosyl transferase HPRT-1 and ATP citrate lyase ACLY-1 were enriched in subsets of glia and neurons, respectively, consistent with scRNA-seq datasets (**Fig. 7A, B**). The cytochrome c oxidase subunit 6B COX-6B was highest enriched in the distal gonad and intestine (**Fig. 8A**), while scRNA-seq predicted highest expression in the pharynx. The tryptophan 2,3-dioxygenase TDO-2 showed highly enriched or specific protein expression in epidermis, whereas its expression predicted by scRNA seq was broader with modest epidermal enrichment (**Fig. 8A, B**). The hydroxyacyl-CoA dehydrogenase F54C8.1, which we independently isolated from the same pool as COX-6B and TDO-2, showed high enrichment in sperm (**Fig. 8B, C**) whereas it was predicted to be enriched in the intestine based on function but correctly suggested by scRNA-seq as highly enriched in or specific to sperm. We note that the gut granule signal in F54C8.1::mScarlet3 tagged worms was comparable to that observed in untagged N2 worms, indicating that it is background autofluorescence (**Fig. 8D**). The heat shock protein HSP-1 was enriched in the germline while predicted to be ubiquitous and evenly expressed by scRNA-seq (**Fig. 9**). Similarly, the triosephosphate isomerase TPI-1 was observed to be enriched in neurons, which was not consistently predicted from scRNA-seq data (**Fig. 9**). Overall, numerous genes with ubiquitous transcript expression showed tissue-specific enrichment not predicted by transcript levels with a general trend towards underestimation of relative protein levels in the germline.

**Figure 4:**
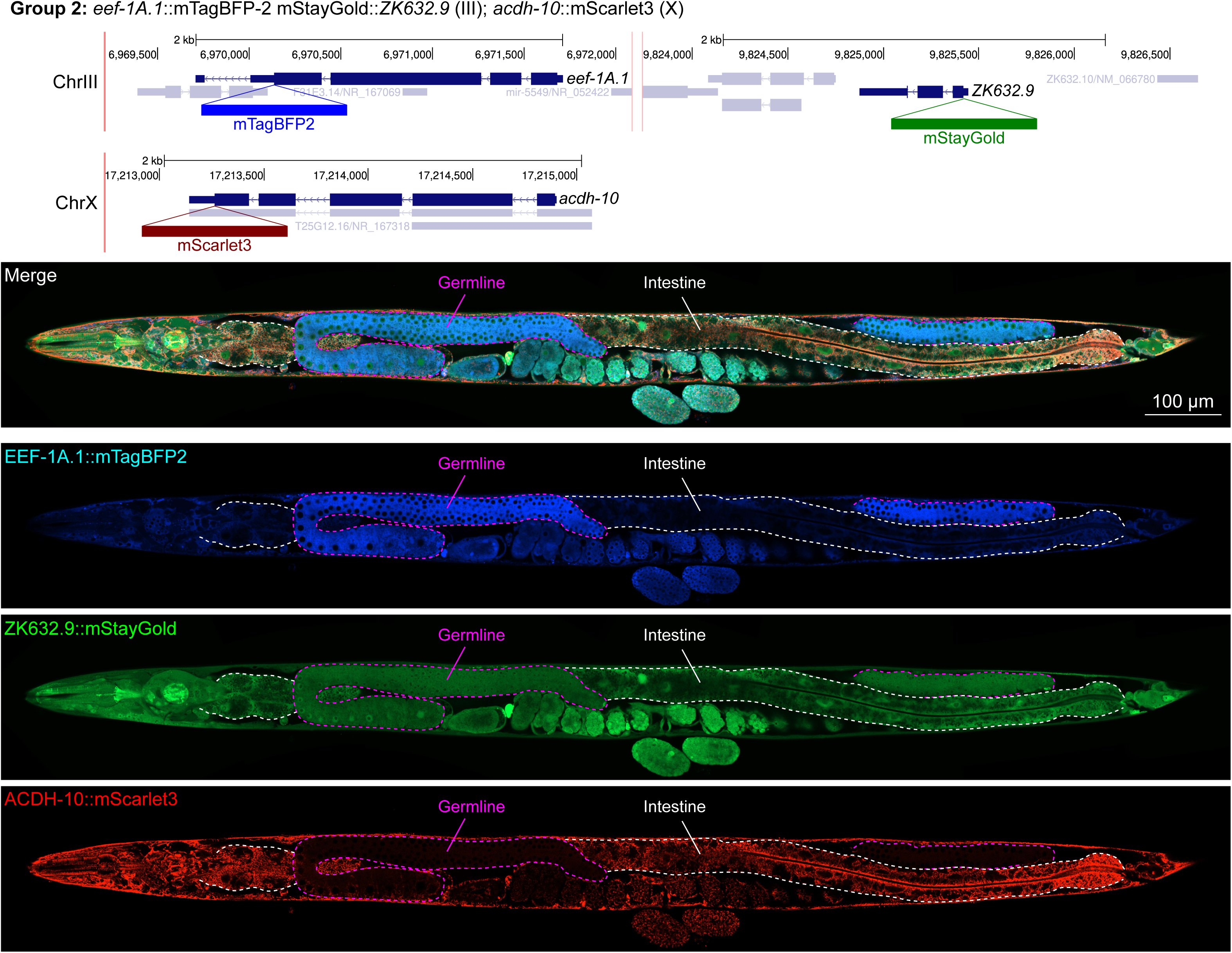
Expression and localization of genes in pool 2. Expressions and localizations of translation elongation factor 1 alpha 1 EEF-1A.1::mTagBFP2, nuclear protein 1 homolog ZK632.9::mStayGold and acyl-CoA dehydrogenase ACDH-10::mScarlet3 proteins in a live adult hermaphrodite worm by confocal microscopy. All three tags were isolated within the same individual then homozygosed at the F2 generation before imaging. Germline and intestine of the animal are delineated in magenta and white dashed lines, respectively. The worm was imaged in separate overlapping frames with high magnification then stitched to reconstruct the whole animal. A single plane near the mid-plane of the animal was imaged.

**Figure 5:**
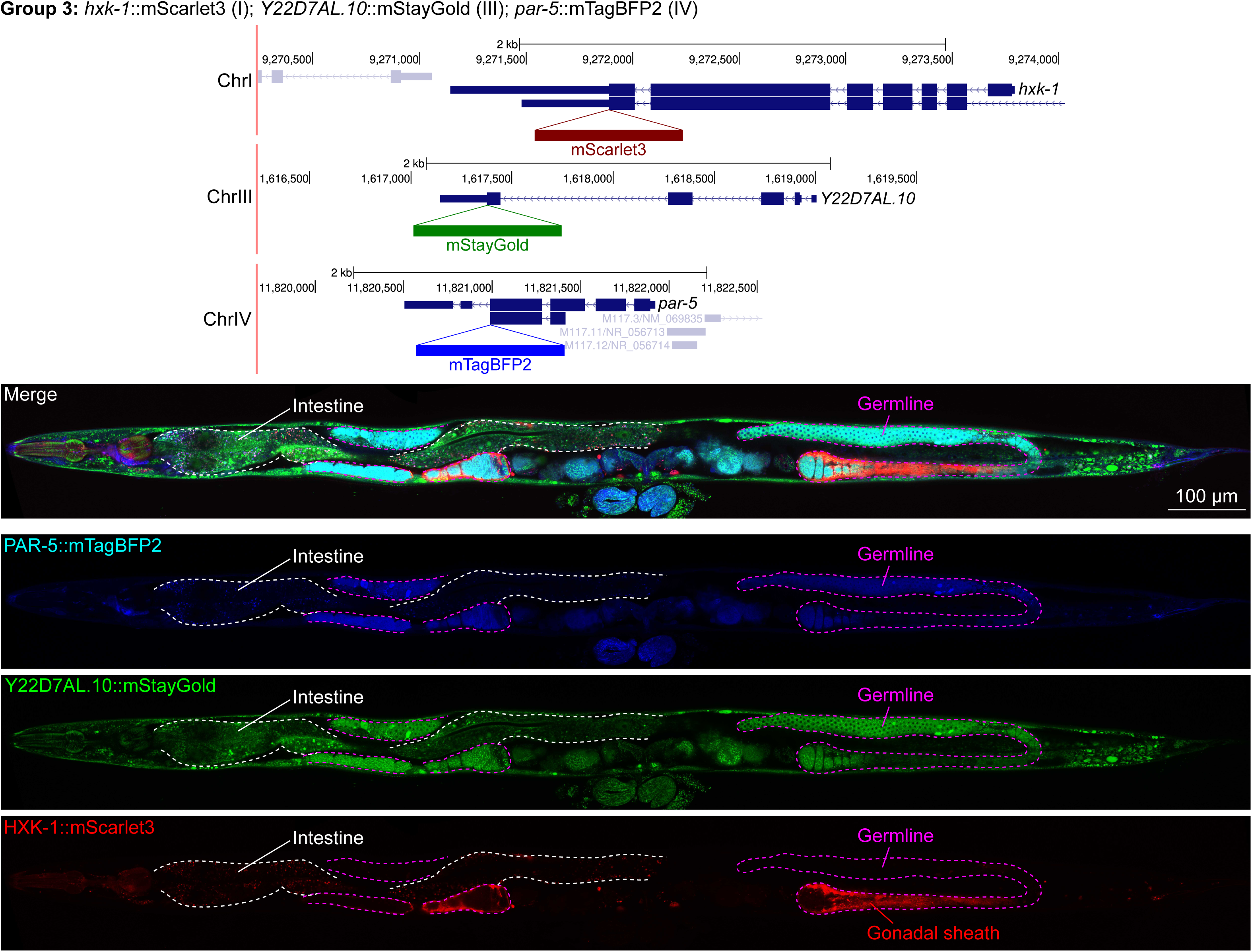
Expression and localization of genes in pool 3. Expressions and localizations of 14-3-3 protein PAR-5::mTagBFP2, heat shock protein family E Y22D7AL.10::mStayGold and the glucokinase HXK-1::mScarlet3 proteins in a live adult hermaphrodite worm by confocal microscopy. All three tags were isolated within the same individual then homozygosed at the F2 generation before imaging. Magenta and white dashed lines, germline and intestine respectively. Image reconstructed from multiple overlapping high magnification images. Single plane near the mid-plane of the animal.

**Figure 6:**
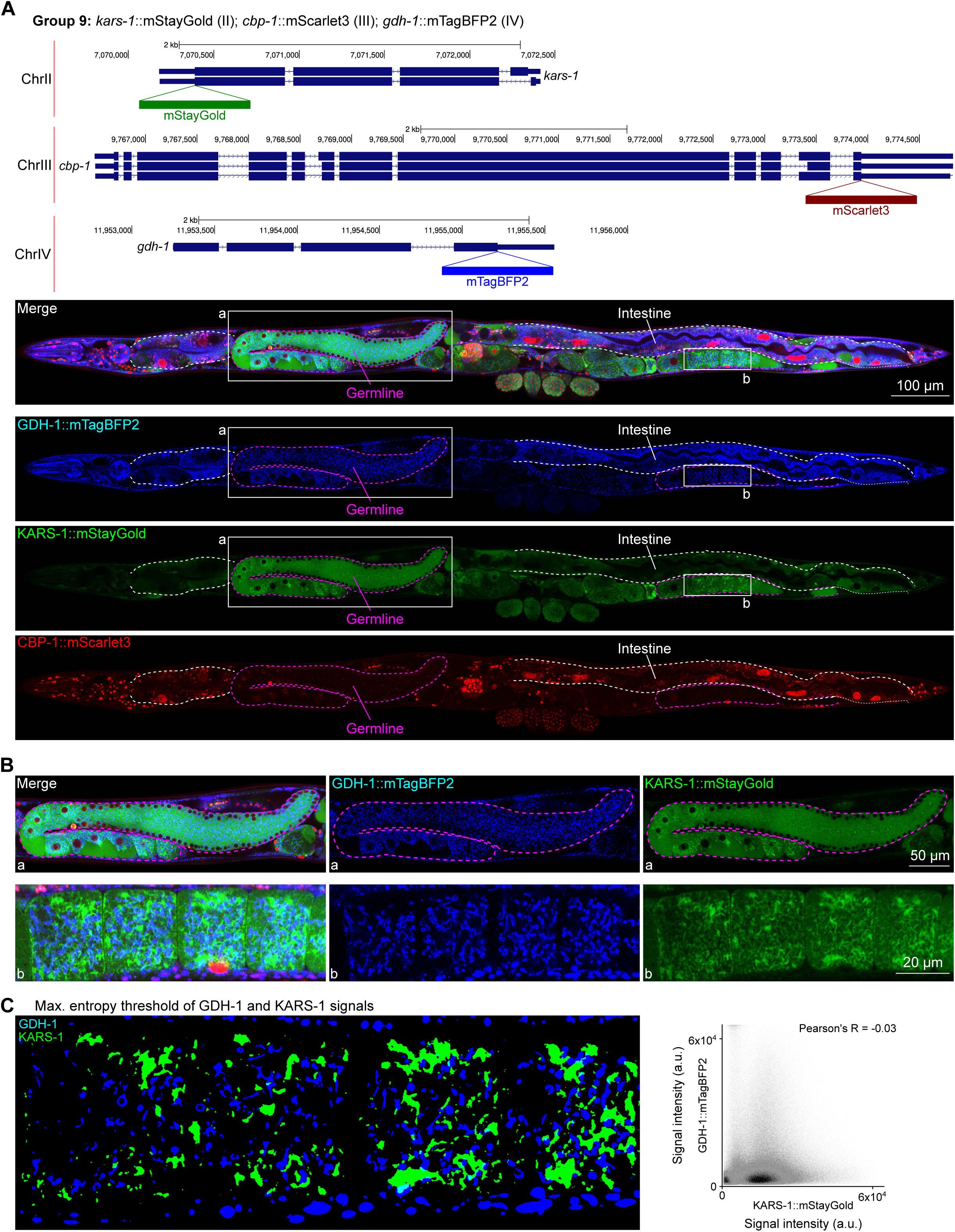
Expression and localization of genes in pool 9. **(A)** Expressions and localizations of glutamate dehydrogenase GDH-1::mTagBFP2, lysyl-tRNA synthetase KARS-1::mStayGold and the CREB binding protein CBP-1::mScarlet3 proteins in a live adult hermaphrodite worm by confocal microscopy. All three tags were isolated within the same individual then homozygosed at the F2 generation before imaging. Magenta and white dashed lines, germline and intestine respectively. Image reconstructed from multiple overlapping high magnification images. Single plane near the mid-plane of the animal. **(B)** Germline insets showing non-overlapping localizations of GDH-1 and KARS-1, predicted mitochondrial proteins. **(C)** Left, maximum entropy thresholds of GDH-1 and KARS-1 signals. Right, correlation plot of signal intensities of GDH-1 and KARS-1.

**Figure 7:**
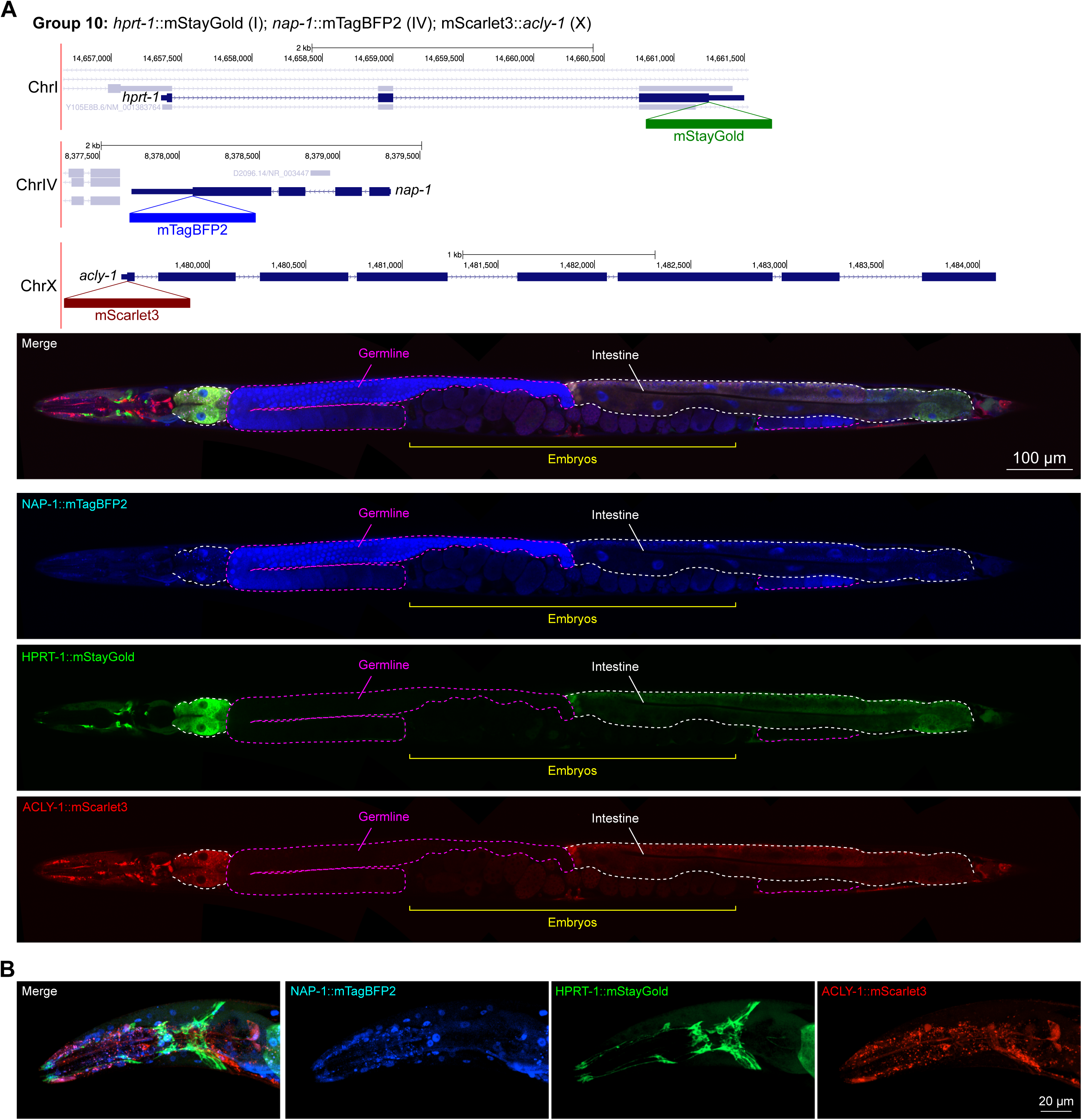
Expression and localization of genes in pool 10. **(A)** Expressions and localizations of nucleosome assembly protein NAP-1::mTagBFP2, phosphoribosyl transferase HPRT-1::mStayGold, and ATP citrate lyase ACLY-1::mScarlet3 proteins in a live adult hermaphrodite worm by confocal microscopy. Magenta and white dashed lines, germline and intestine respectively. Image reconstructed from multiple overlapping high magnification images. Single plane near the mid-plane of the animal. **(B)** Expressions and localizations of nucleosome assembly protein NAP-1::mTagBFP2, phosphoribosyl transferase HPRT-1::mStayGold, and ATP citrate lyase ACLY-1::mScarlet3 proteins in the head of a live adult hermaphrodite worm by confocal microscopy. Image shows a maximum intensity projection of a Z-stack optically sectioned at 1 µm per stack.

**Figure 8:**
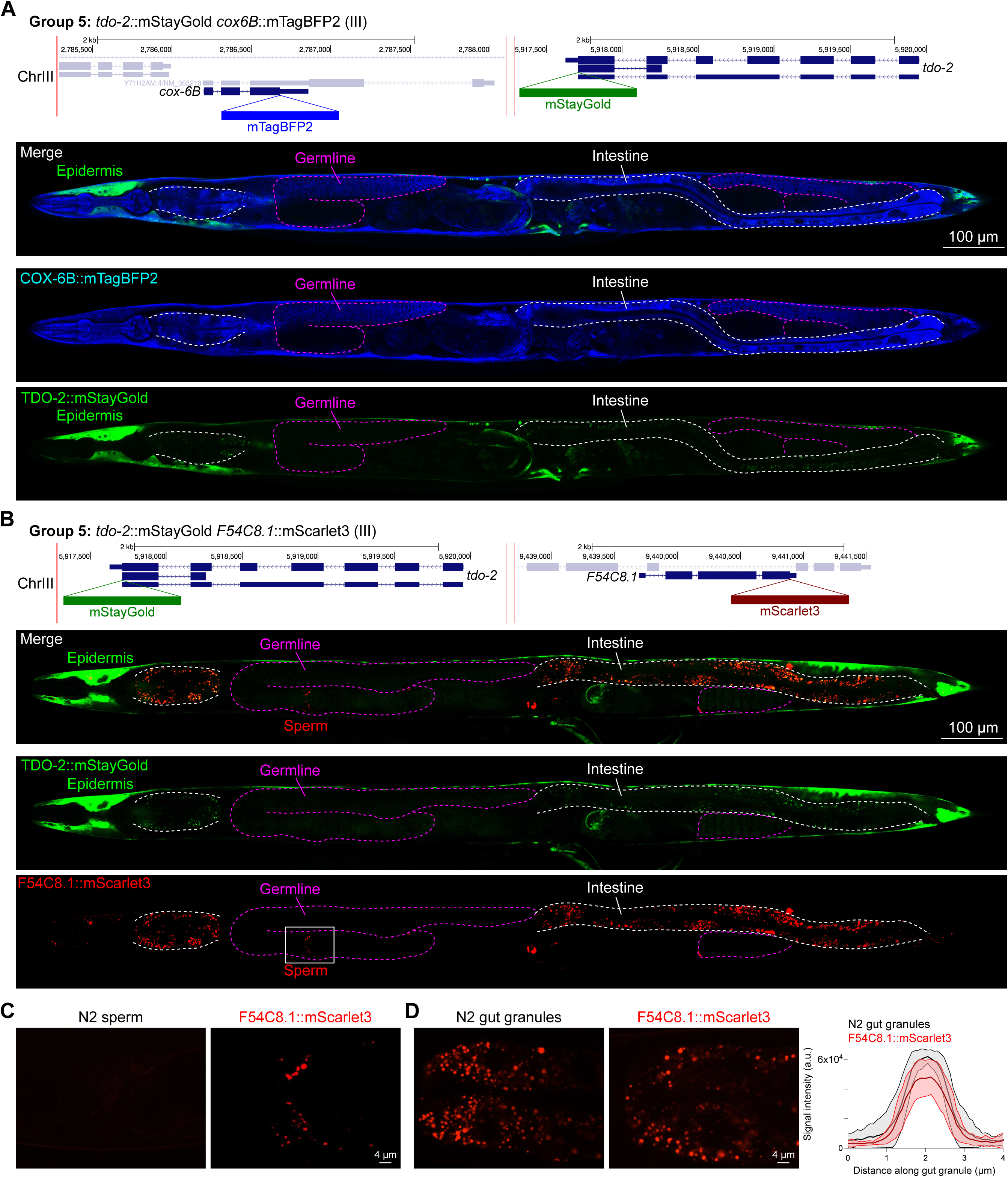
Expression and localization of genes in pool 5. **(A)** Expressions and localizations of cytochrome C oxidase subunit 6B COX-6B::mTagBFP2 and TDO-2::mStayGold proteins in a live adult hermaphrodite worm by confocal microscopy. The two tags were isolated within the same individual then homozygosed at the F2 generation before imaging. Magenta and white dashed lines, germline and intestine respectively. Image reconstructed from multiple overlapping high magnification images. Single plane near the mid-plane of the animal. **(B)** Expressions and localizations of tryptophan 2,3-dioxygenase TDO-2::mStayGold and hydroxyacyl-CoA dehydrogenase F54C8.1::mScarlet3 proteins in a live adult hermaphrodite worm by confocal microscopy. The two tags were isolated within the same individual then homozygosed at the F2 generation before imaging. Magenta and white dashed lines, germline and intestine respectively. Image reconstructed from multiple overlapping high magnification images. Single plane near the mid-plane of the animal. **(C)** F54C8.1::mScarlet3 signal in sperm compared to untagged N2 worms imaged under the same settings. **(D)** F54C8.1::mScarlet3 signal in gut granules compared to untagged N2 worms imaged under the same settings. N = 20 F54C8.1::mScarlet3 granules and 24 N2 granules quantified from 2 worms for each genotype. Lines, mean ± SD.

**Figure 9:**
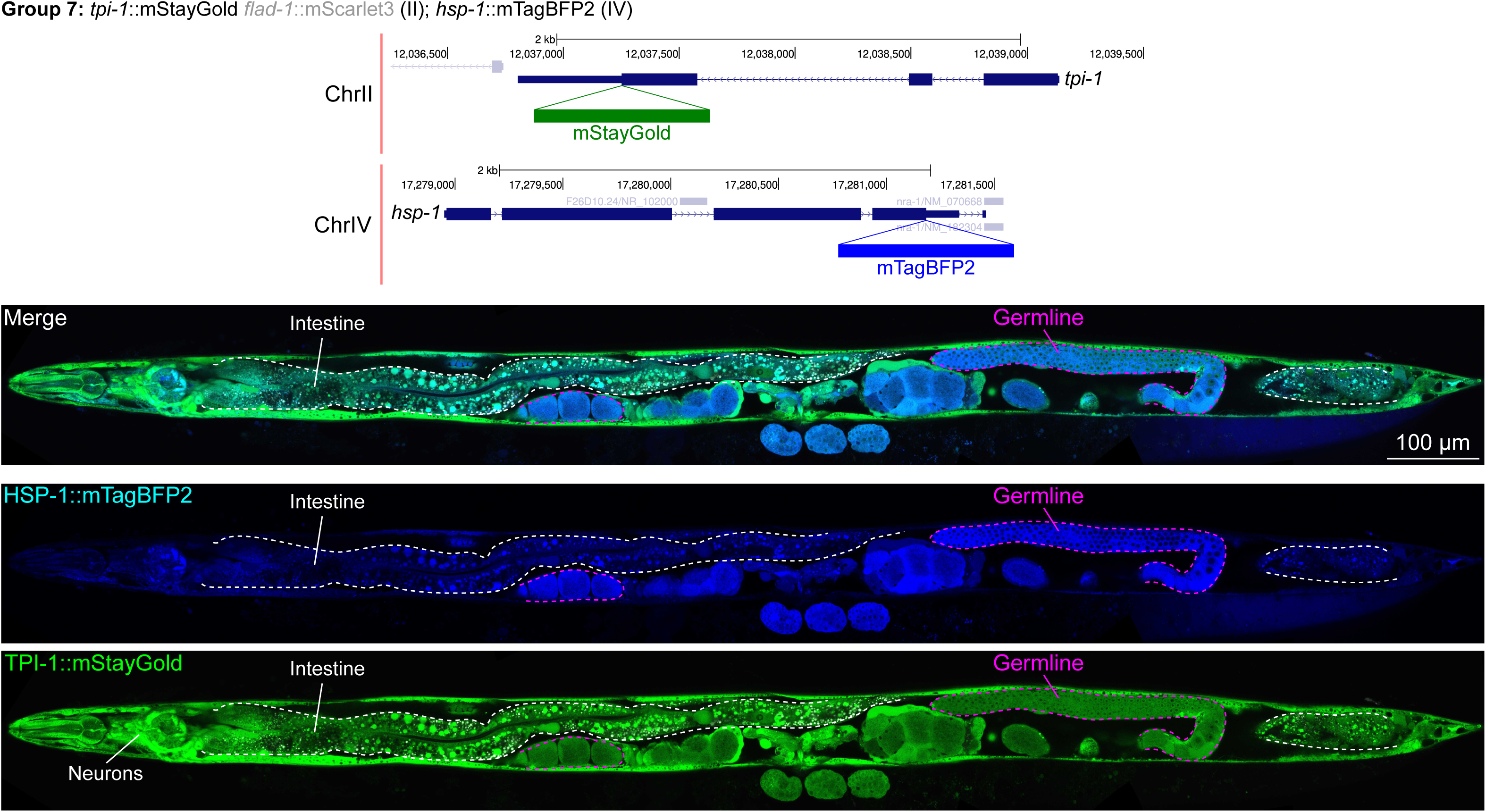
Expression and localization of genes in pool 7. Expressions and localizations of heat shock protein family A HSP-1::mTagBFP2 and triosephosphate isomerase TPI-1::mStayGold proteins in a live adult hermaphrodite worm by confocal microscopy. The two tags were isolated within the same individual then homozygosed at the F2 generation before imaging. Magenta and white dashed lines, germline and intestine respectively. Image reconstructed from multiple overlapping high magnification images. Single plane near the mid-plane of the animal.

### Unanticipated subcellular localization patterns of proteins

Given that our tags were designed to be translationally fused to the proteins of interest, we assessed their subcellular localizations and compared them to localizations annotated experimentally or predicted by homology (Harris et al., 2020). Most notably, we found two proteins, GDH-1 and KARS-1, detected in mitochondria by mass spectrometry (Guo et al., 2025; Li et al., 2009), and co-tagged in the same worm, to display mitochondrial-like localizations; however, they did not overlap (**Fig. 6B, C**). This finding suggests that subsets of mitochondria in the gonad may have varying markers and metabolic specializations. Indeed, a study published around the time of our pilot experiment described metabolically distinct subpopulations of mitochondria depending on cellular energy demand (Ryu et al., 2024).

Furthermore, we found that the V-ATPase VHA-2, involved in proton transport and predicted to be localized to cell and vesicular membranes and detected in membrane fractions by mass spectrometry (Mawuenyega et al., 2003), displayed perinuclear localization in the gonad specifically (**Fig. 10**). This indicates that cell type-specific nuclear envelopes or endoplasmic reticula may have unique demands for pH control which are yet unexplored, to our knowledge.

**Figure 10:**
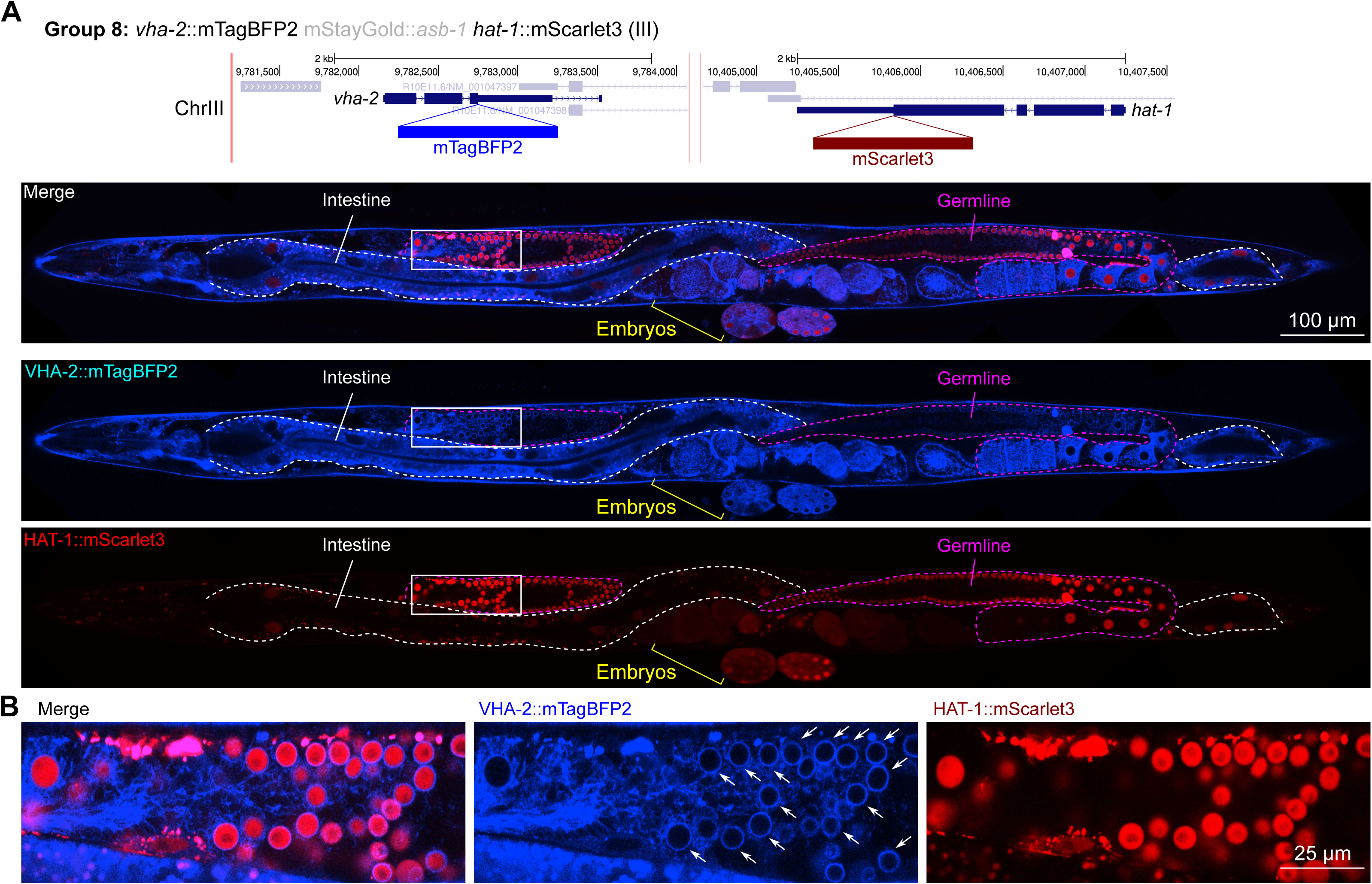
Expression and localization of genes in pool 8. **(A)** Expressions and localizations of ATPase H+ transporting V0 subunit C VHA-2::mTagBFP2 and histone acetyltransferase HAT-1::mScarlet3 proteins in a live adult hermaphrodite worm by confocal microscopy. The two tags were isolated within the same individual then homozygosed at the F2 generation before imaging. Magenta and white dashed lines, germline and intestine respectively. Image reconstructed from multiple overlapping high magnification images. Single plane near the mid-plane of the animal. **(B)** Inset showing germ cell nuclei. The proton pump VHA-2 localizes to nuclear membranes in germ cells specifically, indicated by white arrows.

In general, subcellular localizations were predicted more accurately than cell type-specific abundance, though we observed additional dynamic localization patterns unanticipated by molecular function. For instance, the DNA-binding nuclear phosphoprotein p8 homolog ZK632.9 was predicted to be nuclear but observed diffusely with modest nuclear enrichment (**Fig. 4**). We note that this may be due to its small size of 62 amino acids, which could be influenced by the larger 226 amino acid mStayGold tag. However, the tagged worm does not show any annotated ZK632.9 depletion phenotypes such as sterility (Simmer et al., 2003), indicating that it is at least to some extent functional. Additionally, the histone acetyltransferase 1 HAT-1 showed broadly nuclear localization as predicted but was cytoplasmic in early embryos (**Fig. 10**), indicating dynamic subcellular localization. Finally, the tryptophan 2,3-dioxygenase TDO-2 with previously unknown localization appeared cytoplasmic (**Fig. 8A, B**).

## Discussion

We have provided proof of concept for the ease and efficiency of tagging at least three proteins at once during a single round of injection. This approach also provides the flexibility of tags being isolated together or individually, if desired. We also find that modern-day fluorophores allow the visualization of proteins at a wide range of expression levels, even at the lowest expression levels tested. Extrapolating our results to annotated transcript levels of *C. elegans* genes, we predict that at least 324 proteins could be directly visualized by mTagBFP2, 1578 with mStayGold, and 7249 with mScarlet3 for high throughput screening using low powered fluorescence stereomicroscopy. At minimum, this could enable annotation of one third of *C. elegans* proteins and at best provide sufficient markers to co-tag the entire genome in groups of 3 genes. In the latter scenario, brighter genes would be used as co-CRISPR markers grouped with dimmer genes which would be isolated among bright gene positive worms through alternative screening methods such as PCR or higher magnification, longer exposure fluorescence microscopy (**Fig. 1A**). Given many genes are not broadly expressed, their bulk transcript levels in whole worms are likely to underestimate expression within subsets of cells, which may aid in their detectability and isolation. For instance, we could clearly visualize F54C8.1::mScarlet3 in adult sperm by fluorescence stereomicroscopy despite a bulk FPKM of 16. Similarly, nuclear localized proteins will likely be easier to detect even at low expression levels, given the concentration of signal in small subcellular compartments. Indeed, this helped us detect HAT-1::mScarlet3 (56 bulk FPKM), which may have been too dim if distributed more broadly within cells. Even with the relatively small number of targets tested, we have observed numerous unanticipated patterns of tissue-specific enrichment and subcellular localization.

By tagging essential genes and in most cases observing no overt effect on viability of the animal, we have also shown that many N- or C-terminal tags do not disrupt protein function in an obvious way. Other groups targeting more specific protein classes (e.g., collagens) also observed that most tagged proteins retained function (Ragle et al., 2025). We interpret failed tagging attempts as disruptions to protein function, which would necessarily prevent isolation of viable worms, which occurred in 20% of attempted tags. In our experience, inclusion of an 18 amino acid linker reduces the chances of disrupting protein function, and for all our tags we have included a universal 18 amino acid linker. However, we do fully recognize that N- or C-terminal tagging will not always produce properly localized and/or functional protein. We have shown a few examples of functional disruptions here, and it is well appreciated in the field that many types of proteins are difficult to tag without either disruption of function or localization, or both. We also recognize that in some cases where worms appear superficially wild type, tags may have more subtle effects on protein function which may impair fitness. One strategy to predict whether a fused protein may lose function is by comparing the predicted structure of the fusion to the original protein of interest with Alphafold (Jumper et al., 2021). However, even if the fused protein is predicted to fold normally, steric interactions, multimeric complex formation and other intrinsic properties of the protein such as solubility may still be affected and are difficult to predict based solely on structure. Smaller tags, including epitope tags like 3xFLAG, V5, and SPOT, can be expected to minimize the risk of such disruptions. However, these present their own problems. Classic epitope tags require fixation of samples, thereby preventing live imaging and introducing caveats of tissue deformation during fixation and permeabilization. The Alfa tag system that operates through stabilizing fluorescent protein-fused nanobodies (Götzke et al., 2019) may, for steric reasons, carry the same risk of disruption of endogenous protein function or subcellular localization as covalent tagging. For genes with no obvious mutant phenotypes, assessing the functionality of tagged alleles will be difficult. For such genes, if an interaction-based phenotype such as synthetic lethality is annotated, the tagged gene can be crossed to the interactor to assess synthetic phenotypes. We envision that any novel or unexpected subcellular localizations observed will require validation by independent methods, as in every high-throughput experiment. Overall, the balance between throughput and functionality is, in our view, best struck with translational fusions of fluorophores to proteins of interest, which allows direct live visualization while carrying well understood limitations and artefacts.

Proteins localized to very specific subcellular sites, such as a handful of synapses in the nerve ring, pose significant challenges for accurate cell identification. Moreover, around 17% of the *C. elegans* genome (3,484 genes) may encode for secreted proteins (Suh and Hutter, 2012). Endogenous tagging of a substantial fraction of these proteins could reveal spatial patterns of secretion, distinguishing components that remain near their cell of origin from those that disperse to distal sites (Keeley et al., 2020). Tagging secreted proteins can also reveal sites of secretion – such as apical or basolateral membranes, or neurites – as has been observed for specific insulins (Sural et al., 2025) and for neuropeptides that localize selectively to synaptic regions (Hu et al., 2011; Toker et al., 2025; Tomioka et al., 2006). In cases where the cellular origin of either highly localized or secreted proteins may be difficult to infer, the labeling of the entire cytoplasm of cells (or, alternatively, its membrane) would be necessary. In theory this could be achieved by using a double fluorophore tag, where the second fluorophore is separated by a splice leader or T2A sequence from the protein-tagging, first fluorophore. However, in practical terms, such a cassette is much larger than 1 kilobases, and in our experience larger inserts lead to a significant drop in the efficiency of successful insertion, therefore negatively affecting throughput. Where such cases are observed, a small, self-cleaving T2A peptide sequence, which frees up the tag from the fused protein, can be easily inserted in a second round of CRISPR/Cas9 to validate the cell of origin.

Fluorescent landmarks can facilitate delineation of cell boundaries, nuclei, cytoskeleton, or other organelles like mitochondria. Use of visible fluorophores such as mTagBFP2, mStayGold and mScarlet3 leaves the far-red spectrum open for the inclusion of markers such as membrane-targeted far-red fluorescent proteins like miRFP680 (Zhang et al., 2023) or cell-permeant near-infrared dyes. In an ideal case, pools of target-specific mTagBFP2, mStayGold and mScarlet3 would be directly injected into integrated marker strains expressing miRFP680 targeted to landmarks of interest. Another option is to group genes strategically to include one ubiquitously expressed predicted cytoplasmic gene. With confocal or super-resolution microscopy, such impromptu landmarks may provide some indication of subcellular localization for particularly difficult proteins as a starting point.

Illustrating the discovery aspects of protein tagging, we found that two predicted mitochondrial proteins, KARS-1 and GDH-1, localized to distinct subsets of mitochondria. This is consistent with parallel reports of functionally differentiated mitochondria in cells with varying metabolic demands (Ryu et al., 2024). To increase the chance of such discoveries, it may be informative to group multiple proteins predicted to localize to the same subcellular compartments together in a 3-color co-injection step. With sufficient coverage, this can yield information on the differentiation or uniformity of organelles within the same cell.

We also noted unexpected cell type dependent distributions of proteins involved in broadly important metabolic processes such as ACDH-10, which was depleted from the germline compared to other tissues, and HXK-1, which was highly enriched in the gonadal sheath. Notably, for these as well as other cases, scRNA-seq datasets were not sufficient to deduce *a priori* the observed cell type specific differences at the protein level. Importantly, many genes encoding metabolic enzymes including *acdh-10* and *hxk-1* have paralogs that likely perform similar catalytic functions. Yet, duplicate genes with identical functions are generally not evolutionarily stable (Adler et al., 2014; Lynch and Conery, 2000); thus such genes are likely to differ in some meaningful parameter (e.g., regulation or activity) that might align with tissue-specific functional needs. Fully annotating the expression patterns of paralogs at the protein level could indicate which tissues require unique metabolic needs and indicate which paralogous genes have undergone sub-versus neo-functionalization. For those proteins that are less functionally understood, unexpected distributions might indicate which merit further study.

We routinely inject 30 animals per day, with 5 worms per injection mix. In theory, this can enable injection of 6 pools targeting 18 genes per day, or 90 per week. A single lab can thus expect 1x coverage of the genome in approximately 290 weeks or 5.6 years, working at a modest pace, assuming that 30% of injections will need to be repeated. With simple quality control steps and careful template preparation, such as ensuring the purity of repair templates, we project that fewer injections will need to be repeated. Notably, we find that previously untested guide RNAs and homology arms perform exceptionally well at novel loci, as we only tested one set of reagents for each locus which yielded satisfactory tagging rates. Recent advances in automated injection systems (Pan et al., 2024) combined with fluorescent worm sorters may further accelerate coverage.

The market cost of ordering crRNAs and annealing with tracrRNA is currently $76 USD; the cost of each target-specific primer pair for this study was $23 USD. Approximately $8 USD of commercially purified Cas9 is used per injection mix; $1.53 USD for spin columns. Without the one-time small cost of universal fluorophore template synthesis, the material cost of tagging each gene is $109 USD and would be $2.2 million USD for the entire genome. While the time, material and personnel costs would be daunting for an individual lab, it would be feasible for a group of committed labs within a few years. A centrally coordinated, but distributed effort would avoid wasteful duplications in gene tagging efforts that are already documented in the literature, such as PGL-1, DAF-16 and several Argonaute genes which each have been tagged at least 4 independent times with GFP (Leyhr et al., 2025). The available Caenorhabditis Genetics Center (CGC) strain repository funded by NIH Office of Research Infrastructure Programs (P40 OD010440) as well as emerging database infrastructure dedicated to annotating endogenously tagged strains (Leyhr et al., 2025) will facilitate strain as well as reagent (e.g., guide RNA sequence) cataloging and dissemination.

## Materials and methods

### Nematode maintenance

All *C. elegans* strains were maintained at 20 °C on nematode growth medium (NGM) agar plates with OP50 Escherichia coli as a food source (Brenner, 1974). All CRISPR/Cas9 insertion strains were generated in the Bristol N2 background.

### CRISPR/Cas9-mediated genomic insertions

Sequences of oligonucleotides, fluorophore templates and CRISPR RNAs (crRNAs) are provided in **Supplementary Table 1**. Cas9 protein was ordered from Integrated DNA Technologies (Park Coralville, IA, USA) (Alt-R® S.p. Cas9 Nuclease V3, 500 μg, Cat: 1081059), aliquoted at 5 mg/mL concentration, and stored at -80°C until use. tracrRNA was ordered from IDT (Alt-R® CRISPR-Cas9 tracrRNA, 20 nmol, Cat: 1072532), resuspended in the provided nuclease-free duplex buffer to 0.4 µg/µl (18 µM), aliquoted, and stored at -80°C until use (Ghanta and Mello, 2020).

All guide RNAs were designed using IDT’s guide RNA design tool. The highest ‘on-target’ scoring guide within 10 bp of the desired insert site was selected and ordered at 2 nmol scale as a custom Alt-R™ crRNA from IDT, resuspended in 20 µL nuclease-free duplex buffer (provided by IDT with tracrRNA order), and stored at 100 µM at -20°C.

Single-stranded DNA repair templates were generated as described **(Eroglu et al., 2023)** with minor modifications. Double-stranded linear DNA templates encoding *C. elegans* codon-optimized mTagBFP2, mStayGold(J), and mScarlet3 were ordered as ‘FragmentGENEs’ from GENEWIZ (South Plainfield, NJ, USA) and used as universal templates. Primers encoding 45 bp gene-specific homology followed by universal priming sequences (Eurofins Genomics, 10 nmol scale) were used to introduce homology arms to universal fluorophore templates. One primer was phosphorylated at the 5’ end to enable strand-specific digestion by lambda exonuclease. Templates were amplified using Q5 polymerase from New England Biolabs (NEB; M0491) with 0.5 ng of template per reaction at an annealing temperature of 70°C and extension time of 30 seconds with 40 cycles in total. For each gene, a single 50 µL reaction was run. After amplification, individual 50 µL reactions were pooled in sets of three as indicated in **Table 1** before proceeding to column purification (PureLink™ PCR Purification Kit, Invitrogen, Cat: K310001). Purified pools of double stranded DNA were digested with lambda exonuclease (NEB, cat: M0262) for 20-30 minutes at 37°C followed by micro-column purification (Monarch® Spin PCR & DNA Cleanup Kit (5 μg)), NEB, Cat: T1130) using a modified protocol described previously (Eroglu et al., 2023). For re-runs of pools 4, 6 and 10 which failed initially, we regenerated the repair templates and ensured that after each column purification, the A260/230 ratio of the purified DNA was ≥2.2 and A260/280 was 1.8 ± 0.05 when measured with a Nanodrop spectrophotometer.

Pooled injection mixes were set up by first annealing the crRNAs with tracrRNA as follows: 3.8 μL of tracrRNA and 0.4 μL each of 3 target-specific crRNAs were mixed, then incubated at 95 °C for 5 min followed by 10 °C for 5 min. The Cas9 RNP complex (Paix et al., 2015) was formed by incubating 2.55 μL of tracr-crRNA annealed guide pools with 0.5 μL of Cas9 for 5 min at room temperature. Following Cas9 RNP formation, 4 μL of pooled single stranded DNA template (≥400 ng/µL) was added. Constituted injection mixes were injected into *C. elegans* gonads using standard practice. Progeny of injected worms were screened for fluorescence under a Leica M205 FCA fluorescence stereomicroscope with a Planapo 1.6x M-series objective, pE 300 white MB light source and the following filters: Filter set ET DAPI BP - M205FA/M165FC for mTagBFP2; Filter set ET YFP - M205FA/M165FC for mStayGold; and Filter set ET mCherry - M205FA/M165FC for mScarlet3. Worms positive for fluorescence were isolated on individual plates to establish strains.

### Confocal microscopy

Worms at the adult stage were immobilized with 50mM sodium azide on 5% agarose pads and imaged on a Zeiss LSM980 laser scanning confocal microscope (C-Apochromat 40x/1.20 water objective) with the 405, 488 and 561nm lasers at variable power settings. Detection was on spectral channels of the 32-channel GaAsP-PMT detector with gain at 800 V for all strains except for group 2 (Fig. 4) which was at 725V. For mTagBFP2, detection wavelengths were set to 411-481 nm; for mStayGold, detection was at 490-552 nm; for mScarlet3, detection was at 570-694 nm. Multitracking with the LSM980 acousto-optic tunable filter (AOTF) was used to excite and detect mTagBFP2 and mScarlet3 simultaneously on the first light track (405 nm and 561 nm) while mStayGold was excited and detected on a second track (488 nm) with switching between tracks line by line. Scanning was bidirectional with 8x line averaging; pinhole was set to 31 µM (1.00 AU for mScarlet3). Worms were imaged at 1x sampling (213.13 µm x 213.13 µm at 2576 x 2576 pixels) then stitched together on ImageJ (FIJI) (Preibisch et al., 2009; Schindelin et al., 2012) to reconstitute the whole worm. Worms were linearized using the straighten function on ImageJ.

### Software and statistical analysis

Genome plots showing gene loci were adapted from the University of California Santa Cruz (UCSC) genome browser (http://genome.ucsc.edu) (Casper et al., 2026) and represent ce11 version of the *Caenorhabditis elegans* genome. Plots were generated and statistical analyses performed on GraphPad Prism version 10.6.1.

## Data availability

Reported strains are available from the authors upon request. Source data for Figs. 3 and 8D are provided in **Supplementary Table 2**.

## Acknowledgements

We thank Chi Chen for support with microinjections, members of the Hobert lab, as well as Stephen Nurrish, Emily Bayer, Luisa Cochella, Geraldine Seydoux, Jean-Louis Bessereau, Hari Shroff, David Sherwood, Jake Leyhr, Brent Derry and Piali Sengupta for comments on the manuscript. Matthew Eroglu is supported by the Leon Levy Foundation and the New York Academy of Sciences through a Leon Levy Neuroscience Scholarship. This work was supported by the Howard Hughes Medical Institute.

---

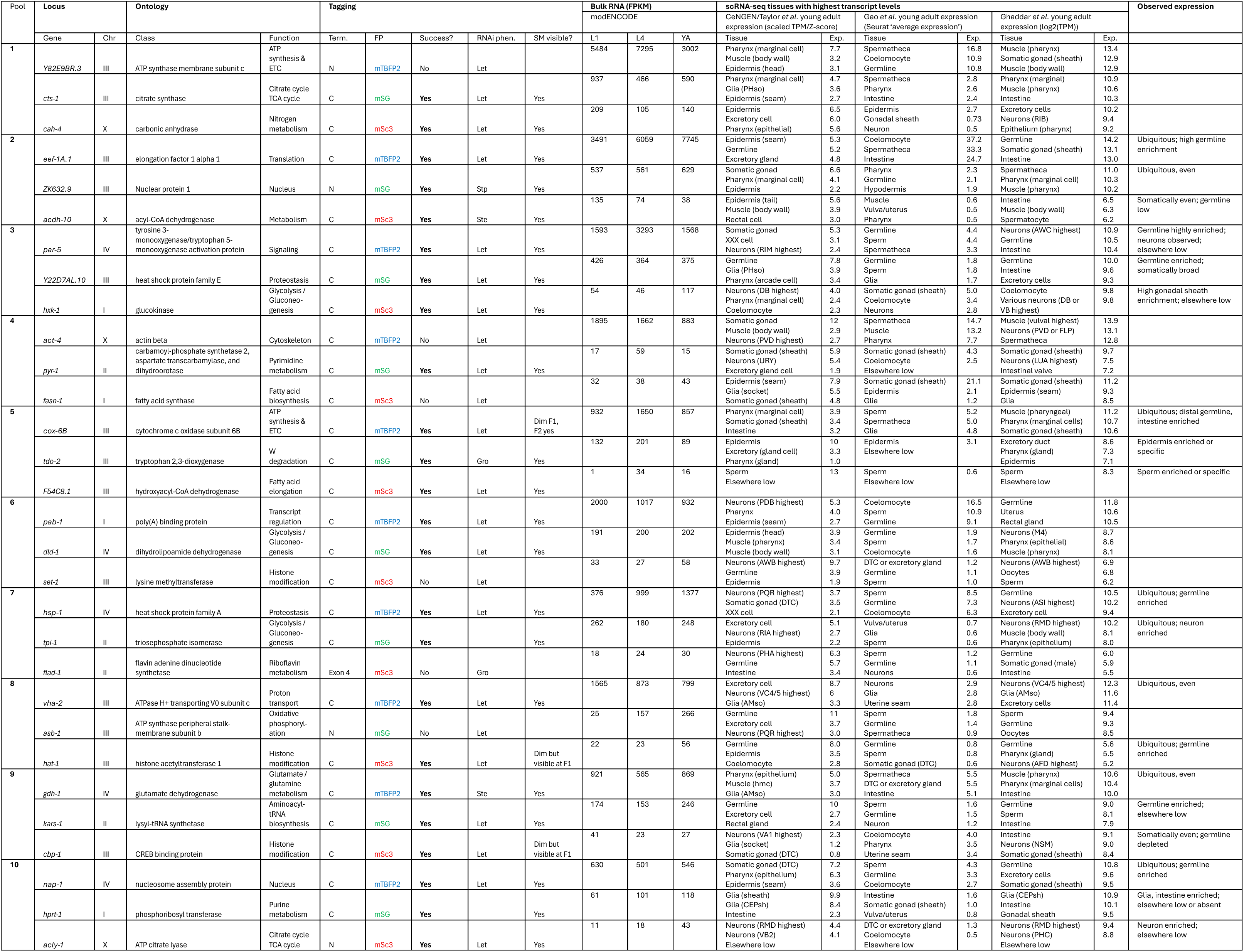

